# Staining-free Visualization of Cell Boundaries in Sliced Tissues for Accurate Binning of Gene Expression in Spatial Transcriptomics

**DOI:** 10.1101/2025.02.22.639623

**Authors:** Zhiming Zhang, Xiaoxia Chen, Youyuan Chen, Zixuan Liao, Tzu-Ming Liu

**Author notes:** Corresponding author: Tzu-Ming Liu. These authors contributed equally.

## Abstract

Overcoming the limitations of sparse expression data in high-resolution spatial transcriptomics remains a major challenge for dimensionality reduction and clustering analysis. Visualizing or inferring tissue cell boundaries using dye staining on the same slice can help assign capture spots on the chip to corresponding unit cells and enhance UMI counts and the number of detected genes. However, the staining process may disrupt tissue structures or degrade mRNA, leading to inaccurate spot assignment and cell annotation. In this study, we employed LysM-Cre mT/mG transgenic mice for spatial transcriptomic analysis, where cell boundaries were delineated by tdTomato signals, and myeloid cells were labeled with enhanced green fluorescent protein (EGFP). With the aid of these fluorescent markers, UMI counts and the number of detected genes per cell exceeded those obtained through naive binning with a square matrix. This staining-free approach for visualizing cell boundaries and landmark proteins provides a valuable framework for the studies of single-cell spatial transcriptomics.

## Introduction

The phenomena of life are governed by tissues and organs composed of diverse cell types. Understanding how cells interact within the tissue microenvironment to perform functions and regulate metabolism is essential for studying organ physiology and disease pathology. According to the central dogma of molecular biology, which describes the flow of genetic information from DNA to RNA to proteins, proteins function as molecular machines that carry out cellular activities, while mRNA transcriptsserve as blueprints, directing protein synthesis based on genetic instructions. Consequently, the transcriptomic expression profile, which comprises expressed genes and their respective expression levels, can be analyzed to infer specific cellular functions and signaling pathways. This, in turn, facilitates the identification/ annotation of cell type, providing insights into the interactions between cellular composition and key molecular factors within the tissue microenvironment. For example, tumor-associated macrophages (TAMs) are critical immune cells within the tumor microenvironment (TME), exhibiting high heterogeneity and playing complex roles in regulating tumor immunity and immunotherapy^[1]^. Analyzing macrophage behavior and designing targeted therapies are considered promising strategies for treating malignant tumors. Therefore, studying novel cell identities in the context is essential for understanding disease progression, ultimately contributing to more precise diagnoses and improved patient outcomes[1].

Compared to the proteome, the transcriptome can be amplified using polymerase chain reaction (PCR), enabling the detection of low-abundance gene transcription activity. Additionally, next-generation sequencing (NGS) has significantly increased the throughput of transcriptomic data analysis, making transcriptomic expression profiling a powerful tool for exploring cellular and molecular activities within tissues. High-throughput transcriptomic analysis via direct RNA extraction from tissues, known as bulk RNA sequencing, has been widely applied in pathological analysis and biological research. However, this technique disrupts cellular structures, preventing the identification of specific cell types from sequencing data. Single-cell RNA sequencing (scRNA-seq) overcomes this limitation by dissociating tissues into individual cells before sequencing their transcriptomes separately, thereby enabling single-cell resolution transcriptomic analysis. However, scRNA-seq is constrained by its requirement for single-cell suspensions, which lead to the loss of spatial information and hinder the localization of specific genes[2], complicating the study of cell-cell interactions and cell-microenvironment dynamics.

Spatial transcriptomics is an innovative technology that deciphers the activity and spatial distribution of genes within a sample, addressing the loss of spatial information inherent in single-cell sequencing[3, 4]. Currently, high-resolution spatial transcriptomics can be broadly classified into two main types: in situ hybridization (ISH) methods and spatial barcoding methods[5]. ISH methods, exemplified by fluorescence *in situ* hybridization (FISH), enable the integration of tissue structure with targeted gene information within the same tissue section at single-cell resolution[6]. However, these methods are limited in their ability to perform whole-genome analysis due to the restricted gene-targeting capacity of probes[7]. Spatial barcoding methods, in contrast, utilize chips embedded with oligonucleotide barcodes to capture mRNA, enabling high-throughput analysis and cell subtype identification[8]. Advances in manufacturing technology have significantly increased the density of capture spots on these chips, allowing spatial transcriptomics to achieve subcellular resolution. This is exemplified by platforms such as 10x Genomics Visium HD (2 µm)[9] and Stereo-seq (500 and 715 nm)[10]. Unlike bulk RNA sequencing, where mRNA is analyzed in bulk tissue extracts, spatial transcriptomics captures mRNA across densely packed capture spots, resulting in a sparse gene-spot matrix[11]. This sparsity poses challenges for cell clustering and annotation due to the low abundance of gene expression at each capture spot.

To address this technical bottleneck, a straightforward approach is to reduce data sparsity by combining RNA expression counts from neighboring spots using a fixed bin size. However, this naive binning method ignores true cell boundaries and mixes transcripts from different cells [9]. A more refined approach involves leveraging cellular imaging data to guide domain segmentation and spot assignment. Hao Chen *et al*. [12] developed Spatial Transcriptomics Cell Segmentation (SCS), which integrates nuclear staining imaging data with sequencing data to infer cell outlines. Similarly, Bohan Zhang *et al*. [13] utilized cell membrane staining to generate more reliable single-cell spatial gene expression profiles. However, these dye-staining-based boundary recognition methods require additional processing steps, which may compromise transcript quality and distort tissue-slice structure. These challenges underscore the need for staining-free cell outline imaging and critical protein landmark visualization to enhance downstream analyses in high-resolution spatial transcriptomics.

Here, we employed LysM-Cre mT/mG transgenic mice^[15, 16]^ to achieve direct visualization of cell boundaries without the need for additional staining. By fusing the myristoylation sites of the MARCKS protein sequence to the N-terminus of full-length tdTomato, the membrane-tagged tdTomato (mT) robustly delineates cell boundaries across various organs. Additionally, membrane-tagged EGFP (mG) was specifically expressed in myeloid cells through LysM-Cre-mediated recombination. Guided by images of these fluorescent markers, the UMI counts and the number of detected genes per cell surpassed those obtained through naive binning with a square matrix. The gene expression of target cells can be analyzed. This study establishes a foundational framework for advancing single-cell-resolved spatial transcriptomics.

## Results

### Two-photon fluorescence imaging of frozen tissue slices from LysM-Cre mT/mG mice reveals clear cell boundaries and myeloid cells’ morphologies

Using two-photon fluorescence imaging, we obtained staining-free images from frozen sections of the liver, heart, and spleen in LysM-Cre mT/mG mice (**Figure 1**). Clear cell boundaries were marked by tdTomato fluorescence (red), while myeloid cells were labeled with EGFP fluorescence (green). In the liver, hepatocytes were distributed throughout the liver lobes, exhibiting prominent tdTomato signals (orange) outlining their membranes, with Kupffer cells (green) interspersed among them. Heart sections, cut longitudinally, revealed cardio myocytes with elongated tdTomato-positive membranes (orange), closely packed together, while cardiac macrophages (green) were observed within the cardiac muscle. In the spleen, splenocytes (orange) appeared smaller in size, with splenic macrophages (green) scattered among them. These results confirm that staining-free tissue slices from LysM-Cre mT/mG mice provide high-quality cell boundary imaging, enabling clear visualization of tissue architecture and myeloid cell distribution.

**Figure 1.**
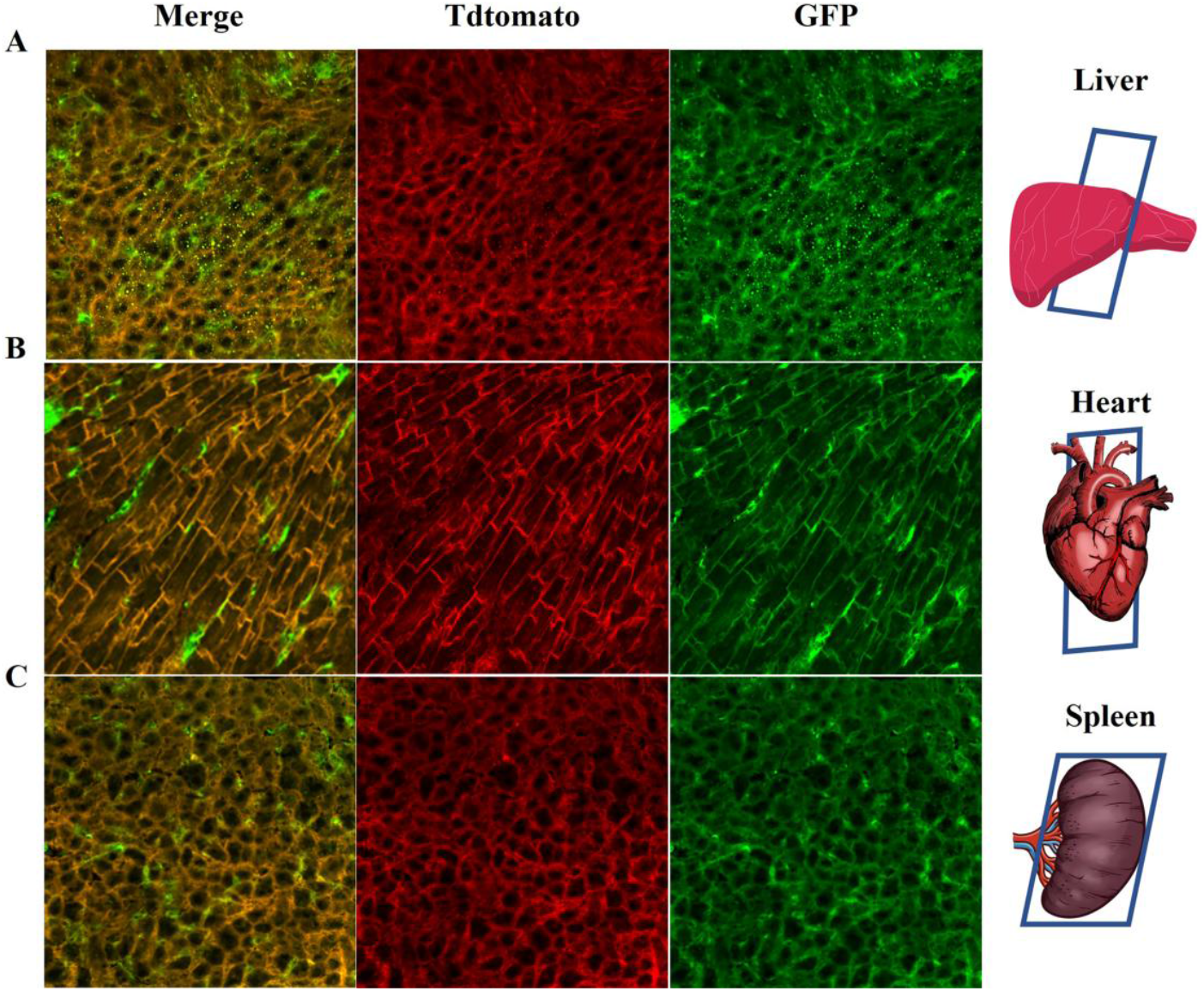
Staining-free two-photon fluorescence images of frozen tissue slice from **(A)** Liver: **(B)** heart: **(C)** Spleen in LysM-Cre mT\mG mice. The tdtomato (red) and EGFP (green) fluorescence were excited at 1040 nm and 960 nm, respectively. Fields of view: 317×317 µm.

### Cell boundary-guided binning of transcripts improves UMI counts/genes per cell in spatial transcriptomics analysis

We performed a high-resolution spatial transcriptomics assay using a high-density spatially barcoded oligonucleotide array embedded on a glass slide (Yosemite, Centrillion Technologies). This array consists of 5000 × 5000 capture spots, each measuring 2 × 2 µm with no gaps, resulting in a 10 × 10 mm capture area. Following large-area two-photon fluorescence imaging, the same liver slice was transferred onto the spatial transcriptomics slide. The RNA integrity number (RIN) of our fresh tissue slices was above 6.5, ensuring high RNA quality for capture. Since tissues are densely packed with proteins such as collagen, laminins, and proteoglycans, which can act as barriers that trap mRNA, we applied an optimized pepsin treatment to partially digest these proteins and release mRNA from the tissue. The tissue slice was then immersed in a hybridization buffer to facilitate mRNA capture. Barcoded oligonucleotides on the slide captured mRNA via their 3′ poly-A tails. Each oligonucleotide contained a unique molecular identifier (UMI) sequence, enabling high-throughput sequencing and precise labeling of captured mRNA molecules. Following reverse transcription, on-chip second-strand synthesis, cDNA amplification, and purification, we obtained high-quality cDNA fragments (300–700 bp). Finally, a sequencing library was prepared for next-generation sequencing (NGS) with a sequencing data volume of 200 G and over 94% sequencing saturation. Raw sequencing data were processed to decode the x-y addresses on the chip and obtain UMI counts per gene using STAR mapping. The address decoding accuracy exceeded 95%, and the gene mapping rate was approximately 70%. The 5000 × 5000 expression matrix was stored in an h5ad file for further analysis.

To align gene expression data with imaging data, we compared the fluorescence image (**Figure 2A**) with the total count reads map (**Figure 2B**) from the same liver slice. Image alignment and registration were performed using structural similarity index measure (SSIM) analysis (**Figure 2C**). Subsequently, cell domains enclosed by tdTomato-labeled membranes were identified and segmented using Cellpose (**Figure 2D**), while EGFP-labeled myeloid cells were segmented using ImageJ (**Figure 2E**). After segmentation, we obtained registered x′-y′ pixel coordinates for each enclosed cell domain. These x′-y′ pixel coordinates (0.31 µm resolution) were then projected onto the x-y feature spot addresses on the gene chip (2 µm resolution) for spatial transcriptomic analysis.

**Figure 2.**
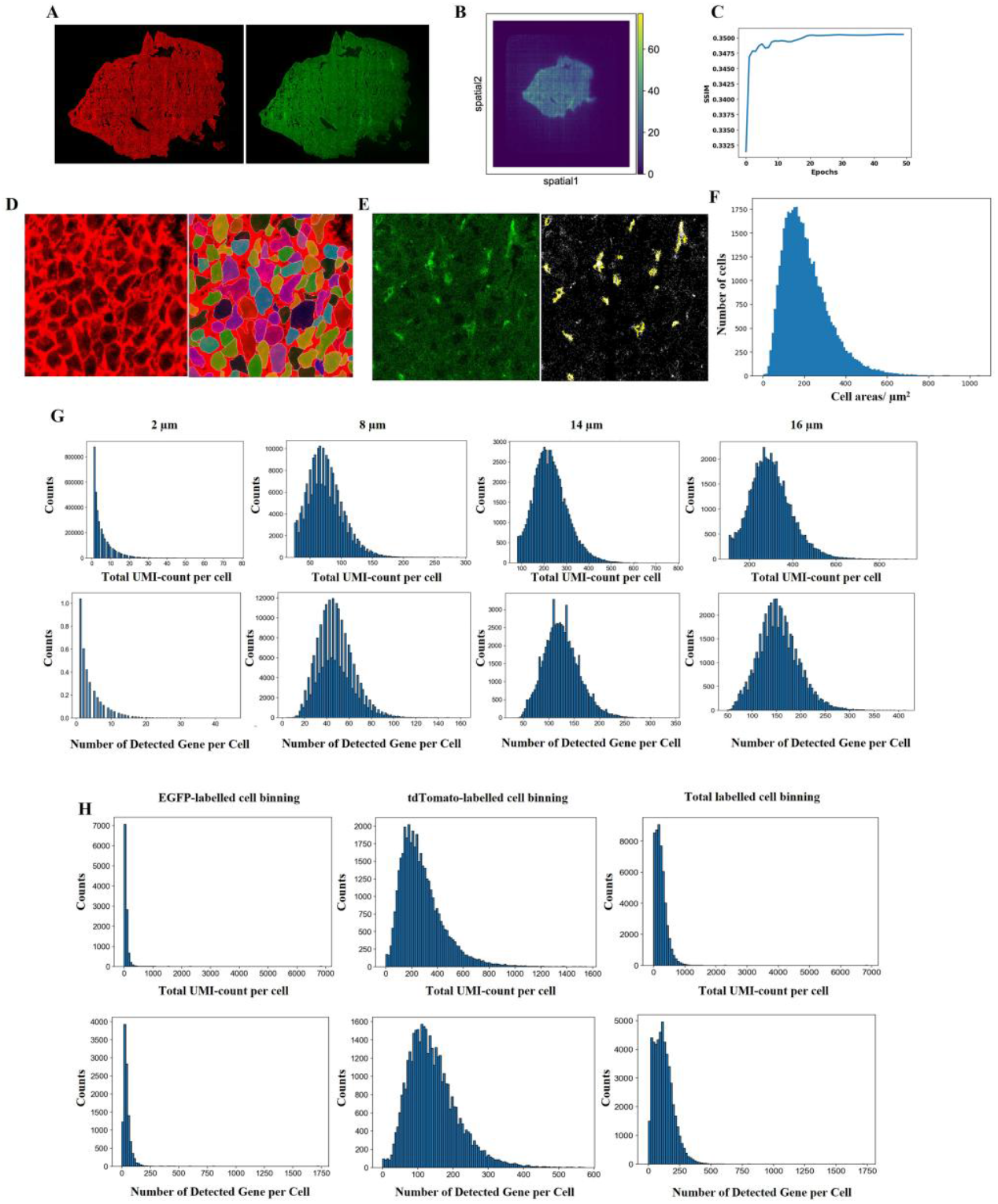
Cell segmentation and binning for spatial transcriptome data. **(A)** The large-scale staining-free fluorescence images of liver slice (red: Tdtomato; green: EGFP). **(B)** The total count reads map of liver slice obtained from spatial transcriptome analysis. **(C)** The SSIM analysis for final registration between fluorescence image and reads map. **(D)** The segmentation of tdTomato-labelled cell membrane (red) in liver slice by Cellpose. Fields of view: 317 × 317 µm. **(E)** The segmentation of EGFP-labelled myeloid cells (Green) in liver slice by ImageJ. Field of view: 317 × 317 µm. **(F)** The area distribution of tdTomato-labelled cells segmented by Cellpose. **(G)** The mean UMI counts/genes per cell at various square bin size. **(H)** The mean UMI counts/genes per cell after cell boundary-guided binning.

Statistical analysis of each 2 µm capture spot revealed low UMI counts and a sparse number of detected genes (**Figure 2G**). This sparsity is a direct consequence of high-density spatial sampling on the chip. To increase gene detection for clustering analysis, a naive square binning approach was commonly used to merge transcripts from neighboring spots. We tested different bin sizes for processing the spatial transcriptome (**Figure 2G**). An 8 µm bin size, commonly used in recent research for single-cell resolution analysis[14], yielded an average UMI count per cell of 79.66 and an average gene number per cell of 51.15—both higher than those obtained under the original 2 µm bin size. The binning significantly reduced the number of zero-count or zero-gene cells and exhibited a well-defined statistical peak in the histogram. To determine the optimal bin size, we calculated the area of Cellpose-segmented cell domains, and the histogram (**Figure 2F**) showed a median area of 187.97µm^2^, corresponding to an optimal bin size of 14 µm. Under this condition, the average UMI count per cell was 228.79 and the average gene number per cell was 125.10. We then applied a 16 µm bin size, which resulted in an average UMI count per cell of 318.64 and an average gene count per cell of 167.44.

After employing cell boundary-guided binning, the results indicated that the mean UMI counts and genes per cell reached 234.50 and 121.77, respectively, which are higher than those obtained using the 8 µm bin size (**Figure 2G**). This suggests that the traditional 8 µm bin size over-segments the cell domain and leads to lower UMI counts per cell. In contrast, the 16 µm bin size overestimates the range of binning and may include mRNA counts from adjacent cell domains. However, we found the results based on cell boundary-guided binning were close to those at 14 µm bin size. Cell boundary-guided binning not only accurately reflect but also enhance the abundancy of mean UMI counts/gene numbers per cell. These findings indicate that staining-free cell boundary-guided binning, enabled by our LysM-Cre mT/mG mice, performs better data quality than naive square binning in spatial transcriptomics analysis.

### The spatial data can perform downstream analysis after convolution and cell boundary-guided binning

In the process of on-chip hybridization, mRNA may diffuse out the cellular boundaries and broaden the gene map of each cell. Considering this factor, we could use convolution function to include mRNA transcript a distance away and enhance gene expression levels before cell boundary-guided binning. For tissue alignment analysis, we first tried a squared 5 × 5 pixels convolution window (roughly 5 µm in radius). Results indicate a significant increase in mean UMI counts and genes per cell (**Figure 3A**). Among the top 20 expressed genes, many genes are closely associated with the liver microenvironment, including the hepatocyte marker gene Alb and genes involved in chemical communication, such as Mup22 and Mup3, which are highly expressed in the mouse liver (**Figure 3B**). Subsequently, we selected three representative gene markers: Alb (hepatocyte marker), Cyp2f2 (periportal zone marker), and Cyp2c29 (pericentral zone marker) to examine their spatial expression levels. The results show that Alb is highly expressed throughout the entire liver slice, while Cyp2f2 and Cyp2c29 exhibit distinct clustering and regional distribution patterns (**Figure 3C**). Therefore, convolution and cell boundary-guided binning not only improve data quality for downstream analysis but also accurately display gene expression information.

**Figure 3.**
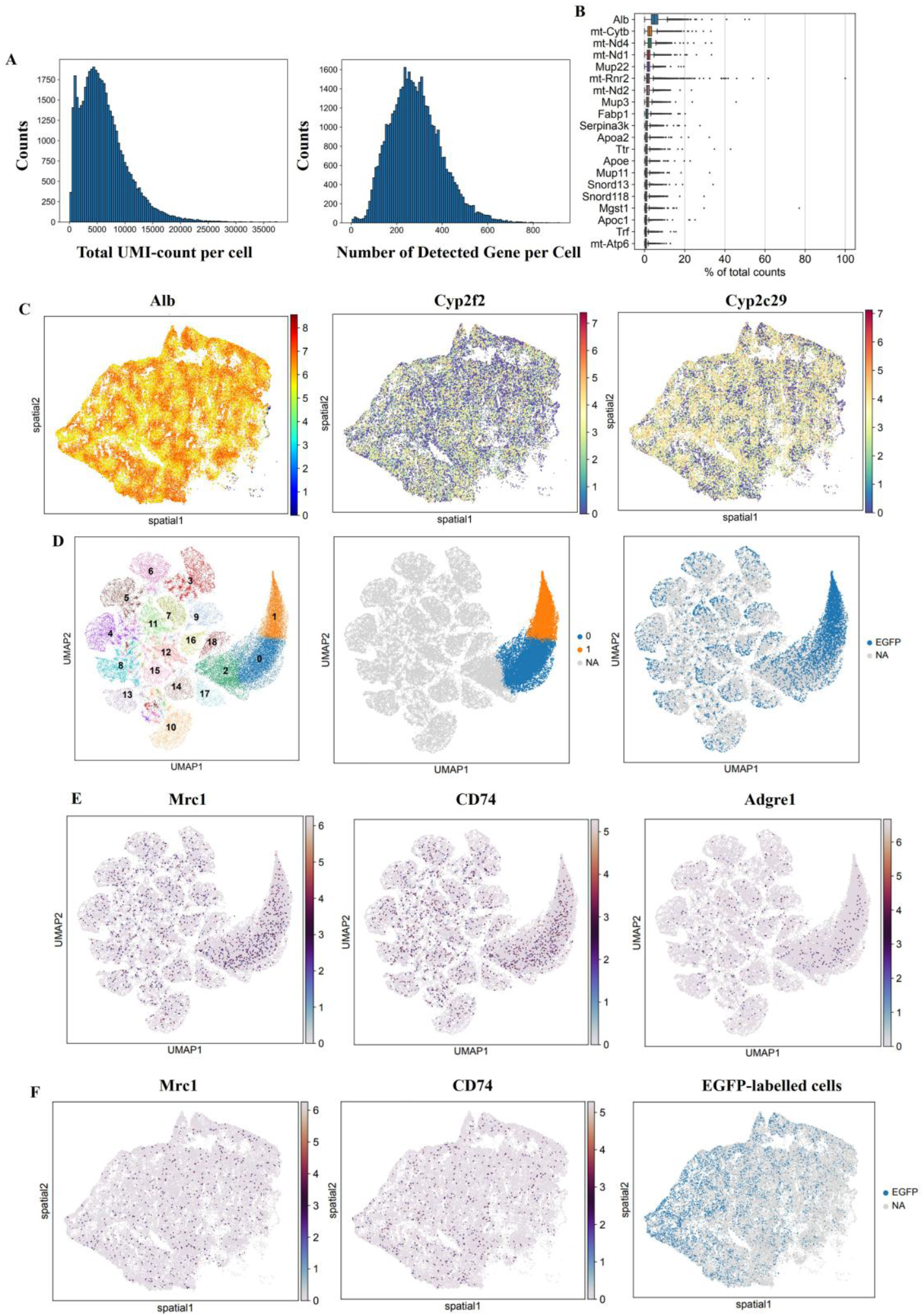
Data analysis after convolution processing. **(A)** The mean UMI counts/genes per cell after convolution and cell boundary-guided binning; **(B)** Top 20 genes in liver sample after convolution and cell boundary-guided binning; **(C)** Alb, Cyp2f2 and Cyp2c29 gene expression in liver slice. **(D)**The results of clustering analysis. **(E)** The expression level of myeloid cell gene markers in various clusters. **(F)** Spatial distribution of myeloid cell marker genes and EGFP-labelled cells.

Subsequently, we performed clustering analysis and identify 19 clusters (**Figure 3D**). We found the EGFP-labelled cells tend to enrich in cluster 0 and 1, and highly expressed myeloid cell marker genes such as Mrc1 and CD74 (**Figure 3D&E**). The results demonstrate the accuracy of clustering analysis. It should be noted that myeloid cell marker gene Adgre 1 do not seem to show obvious high expression in GFP-labelled cells (**Figure 3E**). The possible reason is that the corresponding genes are not expressed at high levels in the tissue itself. Additionally, the myeloid cells marked by Mrc1 and CD74 also show consistency in spatial distribution with the EGFP-labelled cells Overall, these findings indicate that the data possess high quality and accuracy for downstream analysis.

## Discussion

Accurate merging of capture spots from chips is a key step in the analysis of spatial transcriptomics. By utilizing imaging information, precise cell segmentation can guide the merging of capture spots, thereby improving gene abundance for subsequent analysis. Existing cell segmentation methods often rely on complex algorithms or additional staining procedures, which do not fully address the limitations posed by sparse spatial transcriptomic data.

In our project, we employed transgenic mice LysM-Cre mT/mG to obtain staining-free images that provide standardized guidance for cell segmentation. In this type of mice, myeloid cells are also labeled by EGFP, providing spatial landmarks for macrophage annotation. Using a multiphoton imaging system, we first acquired high-resolution images of various tissue slices from the transgenic mice. These slices displayed clear cell boundaries, while landmark signals indicated the locations of myeloid cells. Following imaging, we utilized the same liver slice to perform spatial transcriptomics with the aid of a high-density spatial transcriptomics chip. High-quality data were generated after convolution and cell boundary-guided binning. Overall, our findings suggest that staining-free imaging holds significant promise for guiding cell domain binning in spatial transcriptomics, offering exceptional prospects for understanding the spatiotemporal details of the disease microenvironment.

However, we still need to perform convolution to enhance gene expression levels for downstream analysis due to the sparsity of the data matrix. In the future, we plan to further improve RNA capture efficiency to address this issue. By optimizing permeabilization conditions and enhancing tissue mRNA mobility, we aim to establish novel spatial transcriptomics procedures with high RNA capture efficiency, which will be beneficial for accurate clustering and annotation analysis.

## Materials and Method

### Mouse model and Tissue slice preparation

The transgenic mice Lysm-Cre\mTmG were generated from the hybridization of B6.129P2-Lyz2^tm1(cre)Ifo^/J mice and STOCK Gt(ROSA)26So^rtm4(ACTB-tdTomato,-EGFP)Luo^/J mice, in which myeloid cells express EGFP while other cells express tdTomato [15]. In this experiment, 6-to 8-week-old transgenic mice LysM-Cre\mTmG were utilized. We aimed to determine whether tissue slices from these mice are of sufficient quality for subsequent studies. Various fresh tissue samples, including heart, spleen, and liver, were collected and embedded in optimal cutting temperature compound (OCT). The embedded samples were flash-frozen in liquid nitrogen using isopentane as the medium. Finally, the tissue samples were sectioned using a cryostat (Leica, CM3050S, Germany) and stored at -80 °C prior to use.

### Multiphoton imaging on the tissues slices

Before imaging, slices were fixed with 4 % formaldehyde for 10 min at room temperature. And then the samples were washed with PBS (Phosphate buffered saline) to remove formaldehyde. The slices were immersed with PBS and covered with coverslip. Finally, the slices were observed under Nikon Eclipse Inverted Multiphoton Microscope (A1MP+Eclipse Ti-2E, Nikon instrument Inc., Japan) with a 40×NA=1.15 water-immersion objective. The fluorescence images were obtained under the excitation wavelength of 960 nm and 1040 nm. Under 960 nm excitation, we collect EGFP two-photon fluorescence signals at 506 nm - 593 nm wavelength range. Under 1040 nm excitation, we collect tdTomato fluorescence signals at 604 nm - 678 nm wavelength range. The laser power after the objective was below 20 mW to avoid sample damage. We also acquired large-scale confocal imaging with 19742 × 15020 pixels images at a 2.42 µs pixel dwell time within 1 h under the excitation wavelength of 488 nm and 561 nm. Under 488nm excitation, we collect EGFP fluorescence signals at 500 nm - 550 nm wavelength range. Under 561 nm excitation, we collect tdTomato fluorescence signals at 570 nm - 620 nm wavelength range. The image was composed of 20×15 patches with 5% overlap.

### Liver sample processing for Spatial transcriptomics

After microscopy imaging, the same liver slice was applied for spatial transcriptome based on spatial chip produced by Centrillion Technologies[16]. The related experiment operations were carried out according to product manuals. Briefly, the sample slices were permeabilized with pepsin for 5 min at 37 °C. The sample was inverted onto the chips immersed with hybridization buffer and incubated for 4 h at 42 °C. At this incubation step, mRNA diffused to buttom of chips, where polyT tail can capture polyA cap of mRNA transcript. Amplified cDNA was obtained after reverse transcription, second-strand synthesis and PCR amplification. And then cDNAs with the size of 300-700 bp were collected after magnet beads selection (Backman, B23317). Through fragmentation, end repair, ligation and adapter addition, the library was established with NEBNext Multiplex Oligos for Illumina (NEB, E7335), followed by sequencing using DNBSEQ-T7-PE150 (MGI, China).

### Spatial transcriptome data preprocessing

The raw sequencing data were stored in a FASTQ file, which needs to be transformed into h5ad files for subsequent analysis using Python on Linux system. Initially, Unique Molecular Identifiers (UMIs) and coordinates were extracted from the raw FASTQ files using PostMaster developed by Centrillion Technologies. The reads were then mapped to the reference genome GRCm39 with the STAR aligner. The resulting file was processed by FeatureCounts to generate a count matrix. Subsequently, the Bamreader, also developed by Centrillion Technologies, was used to create CSV files containing the gene ID, UMI, and spatial coordinates. Finally, the resulting file was converted into h5ad format for downstream analysis.

### Spatial registration between captured UMIs and fluorescence imaging

A UMI count matrix of 5000 × 5000 pixels encompassing 22,291 genes was projected into a single image, and its expression levels were normalized to a range of 0–255. For spatial registration, GUI software based on the PyQt5 toolkit was developed to adjust the parameters of image rigid transformation, including rotation and translation. The gene image was initially fixed and then adjusted to match the screen resolution. Subsequently, the fluorescence images were added, also adjusted to match the screen resolution, and rescaled to achieve the same resolution as the gene image. The rescaling factor of 0.155 was determined by the pixel resolution ratio of fluorescence images (0.31 µm) over gene images (2 µm). Following this, the fluorescence images were aligned with the gene image through rotation and translation, with any fluorescence image portions outside the overlapping range being cropped. Relative to the gene image, the fluorescence intensity of the missing fluorescence image part is filled by 0. During this process, an affine transformation matrix that can be defined within the software was applied as follows:

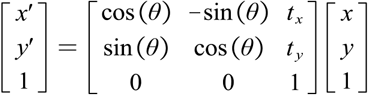

where *P* (*x,y*), *P* (*x*’,*y*’) are the original and affine transformed points. θ is applied to control the rotation of image, and the translation on horizontal and vertical is implemented by *t*_*x*_,*t*_*y*_.

Next, an image similarity evaluation method, SSIM (Structural Similarity Index Measure) was utilized to optimize the best alignment of fluorescence imaging:

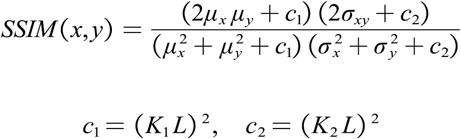

Where *μx,μy* are the mean values of total gene expression image and fluorescence image. 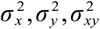 are their variances and covariance. *K*_1_ = 0.01, *K*_2_ = 0.03, *L* = 255 are constant.

### Cell boundary-guided binning on spatial transcriptome

After obtaining aligned fluorescence images, cell boundaries with tdTomato signals were segmented using Cellpose with the cyto2 model. Myeloid cells labeled with EGFP fluorescence signals were identified using ImageJ software, and cells with areas smaller than 60 pixel^2^ were filtered out. To align the coordinates of the fluorescence image with the gene expression data, each pixel of the gene expression image was enlarged into a square on the fluorescence image according to their pixel resolution and the registration information. Finally, pixel coordinates and the center coordinates of cells on the gene chips were used to bin the entire spatial transcriptome. For comparison, different naive square bin sizes were also set to process the data.

### The clustering analysis of spatial transcriptome

To enhance gene abundance for downstream analysis, a convolution operation with a squared 5 × 5 pixel window was applied to process the h5ad files, followed by cell boundary-guided binning of the data. The data was then processed using the Scanpy package on the Python platform for clustering analysis. In brief, the Pearson residuals algorithm, with components set to 15, was employed to identify highly variable genes, from which the top 5,000 genes were selected for dimensionality reduction using PCA (Principal Component Analysis) and the UMAP (Uniform Manifold Approximation and Projection) algorithm. Finally, the Leiden algorithm was applied to perform clustering analysis with a resolution set to 0.3.

## Acknowledgements

This work was supported by University of Macau (File no. MYRG-CRG2022-00009-FHS and MYRG-GRG2023-00053-FHS-UMDF) and the science and technology development fund, Macau SAR (File no. 0002/2021/AKP, 0007/2021/AKP, 0003/2023/RIC, 0013/2023/RIC, 0054/2024/RIB1, and 0038/2024/ITP2).

## Reference

1. Jovic, D., et al., Single-cell RNA sequencing technologies and applications: A brief overview. Clin Transl Med, 2022. 12(3): p. e694.

2. Chung, B.K., et al., Spatial transcriptomics identifies enriched gene expression and cell types in human liver fibrosis. Hepatol Commun, 2022. 6(9): p. 2538–2550.

3. Williams, C.G., et al., An introduction to spatial transcriptomics for biomedical research. Genome Med, 2022. 14(1): p. 68.

4. Rao, A., et al., Exploring tissue architecture using spatial transcriptomics. Nature, 2021. 596(7871): p. 211–220.

5. Cho, C.S., et al., Microscopic examination of spatial transcriptome using Seq-Scope. Cell, 2021. 184(13): p. 3559–3572 e22.

6. Janesick, A., et al., High resolution mapping of the tumor microenvironment using integrated single-cell, spatial and in situ analysis. Nat Commun, 2023. 14(1): p. 8353.

7. Zhou, Y., et al., Encoding Method of Single-cell Spatial Transcriptomics Sequencing. Int J Biol Sci, 2020. 16(14): p. 2663–2674.

8. Du, J., et al., Advances in spatial transcriptomics and related data analysis strategies. J Transl Med, 2023. 21(1): p. 330.

9. Benjamin, K., et al., Multiscale topology classifies cells in subcellular spatial transcriptomics. Nature, 2024. 630(8018): p. 943–949.

10. Wei, X., et al., Single-cell Stereo-seq reveals induced progenitor cells involved in axolotl brain regeneration. Science, 2022. 377(6610): p. eabp9444.

11. Long, Y., et al., Spatially informed clustering, integration, and deconvolution of spatial transcriptomics with GraphST. Nat Commun, 2023. 14(1): p. 1155.

12. Chen, H., D. Li, and Z. Bar-Joseph, SCS: cell segmentation for high-resolution spatial transcriptomics. Nat Methods, 2023. 20(8): p. 1237–1243.

13. Zhang, B., et al., Generating single-cell gene expression profiles for high-resolution spatial transcriptomics based on cell boundary images. GigaByte, 2024. 2024: p. gigabyte110.

14. Polanski, K., et al., Bin2cell reconstructs cells from high resolution Visium HD data. Bioinformatics, 2024. 40(9).

15. Li, Y., et al., Imaging of macrophage mitochondria dynamics in vivo reveals cellular activation phenotype for diagnosis. Theranostics, 2020. 10(7): p. 2897–2917.

16. Ding, X., et al., Spatial Transcriptomics Sequencing of Mouse Liver at 2µm Resolution Using a Novel Spatial DNA Chip. BioRxiv, 2024.

